# Concept-Level Semantic Representations Remain Decodable in Chronic Post-Stroke Aphasia

**DOI:** 10.64898/2026.07.24.740313

**Authors:** Alexander M Swiderski, Jason W Bohland, Jeffrey P Johnson, Michael Walsh Dickey, Stephen M Wilson, William D Hula

## Abstract

Aphasia is characterized by impaired word retrieval, yet most cognitive models of word production assume that underlying conceptual-semantic representations are largely preserved. This study investigated whether concept-level semantic structure remains decodable from BOLD signals in chronic post-stroke aphasia and which semantic models best explain neural representational geometry during covert semantic feature generation. Eight healthy adults and six individuals with chronic aphasia completed a dense-sampling fMRI protocol in which they viewed 57 pictured nouns while silently generating semantic features. Representational similarity analysis showed that an experiential model (Exp48) best matched neural geometry in both people with aphasia and controls, outperforming taxonomic (WordNet) and distributional (Word2Vec, GloVe) models. Using representational similarity decoding, concept identity was recovered well above chance in both groups. No relationship was found between decoding accuracy and language measures from individuals with aphasia. These findings suggest that experiential semantic structure remains robustly represented and decodable in chronic aphasia despite lesion-related language impairments, highlighting preserved conceptual representations alongside altered anatomy.

## Introduction

Aphasia is an acquired language disorder affecting more than 2.4 million people in the United States (Simmons-Mackie & Cherney, 2018). Anomia, or the impaired ability to access and retrieve words, is a hallmark deficit of aphasia (Goodglass & Wingfield, 1997) and a common target of aphasia rehabilitation (Tierney-Hendricks et al., 2021). Cognitive models of word production generally distinguish conceptual-semantic processing from subsequent lexical and phonological access, with semantic information serving as input to word-form retrieval (Dell et al., 1997; Levelt, 1999; Roelofs, 2014; Ueno et al., 2011). Within this framework, naming impairment may reflect degraded conceptual knowledge, impaired access to otherwise preserved semantic representations, disrupted lexical-phonological mapping, or some combination of these mechanisms.

Two central questions arise from this premise. First, which computational model of semantic structure best captures neural representational geometry in people with aphasia? Second, are concept-level semantic representations decodable in chronic post-stroke aphasia? The present study addresses these questions using fMRI, representational similarity analysis (RSA), and representational similarity decoding (RSD) during covert semantic feature generation.

### Semantic models and neural representational geometry

The cognitive models of word production described above specify how conceptual information supports lexical and phonological access, but they are largely agnostic about the underlying representation of that conceptual input. Competing accounts of semantic representation make different predictions about how similarity among concepts is organized. Symbolic or amodal accounts emphasize abstract, category-based relations among concepts (Collins & Loftus, 1975); distributional accounts emphasize statistical regularities in language use (Harris, 1954); and grounded or experiential accounts emphasize perceptual, motor, affective, and contextual dimensions of experience (Barsalou, 2009; Murphy, 2004).

Computational semantic models provide testable instantiations of these accounts. In representational similarity frameworks, each model’s predictions can be expressed as a representational dissimilarity matrix (RDM), allowing the semantic similarity structures to be compared with observed neural similarity structures. These models generally fall into three broad categories – taxonomic, distributional, and experiential – which closely align with the three broad classes described above. Taxonomic models, such as WordNet (Fellbaum, 2010), organize concepts hierarchically into superordinate and subordinate categories and closely align with symbolic or amodal theories (Fodor, 1983). Distributional models such as Word2Vec (Mikolov et al, 2013) and GloVe (Global Vectors for Word Representation; Pennington et al., 2014) capture word meaning through statistical patterns of co-occurrence, representing words that appear in similar contexts (e.g., “cat” and “dog”) as more similar than those occurring in distinct contexts (e.g., “cat” and “shirt”). While not tied to a single theoretical account, distributional models implicitly capture linguistic regularities that may reflect symbolic, grounded, or embodied aspects of meaning depending on context (Lund & Burgess, 1996; Raaijmakers & Shiffrin, 1980). Finally, experiential models such as Experiential 48 (Exp48; Binder et al., 2016), represent concepts using features derived from human experiences, including sensory, motor, affective, and contextual attributes. Exp48 was explicitly designed to reflect functional divisions of the human brain, drawing on evidence that conceptual knowledge is encoded within perceptual, motor, affective, and other modality-specific systems. Its features reflect neurocognitively motivated dimensions of experience rather than abstract or linguistically defined attributes.

Prior neuroimaging work supports the use of these models to test how conceptual information is organized in the brain. Semantic processing engages a distributed, left-dominant network including inferior frontal, temporal, angular, and posterior cingulate regions (see Binder et al., 2009), which serves as a theoretically motivated region of interest (ROI) for the present study. Beyond identifying *where* semantic processing occurs, multivariate methods have begun to investigate *how* conceptual information is represented. Using representational similarity analysis (RSA; Kriegeskorte et al., 2008), Fernandino et al. (2022) demonstrated that experiential models, particularly Exp48, explained unique variance in multivoxel similarity patterns within a semantic network ROI using representational similarity analysis. Tong et al. (2022) further showed that taxonomic, experiential, and distributional models predicted partially distinct subregions of the semantic network using searchlight RSA.

Complementing these RSA findings, prior fMRI work using encoding and decoding models demonstrated that conceptual information evoked during feature generation tasks yields highly structured and predictable patterns of brain activity. Mitchell et al. (2008) showed that whole-brain fMRI activation patterns for concrete nouns could be predicted from semantic feature vectors grounded in sensory–motor co-occurrence statistics for a subset of voxels distributed across the cortex. Anderson et al. (2016) built directly on this foundation by introducing representational similarity decoding (RSD), a method that predicts neural activation for a held-out item by weighting previously observed activation patterns according to their similarity in a model-derived semantic space. Using the same covert feature-generation dataset collected by Mitchell and colleagues (2008), they demonstrated that concept identity could be decoded from the representational geometry of subject-specific BOLD signals. Together, these studies establish that covert semantic feature generation elicits richly structured activation patterns that reflect the underlying organization of conceptual knowledge. In the present study, we adapt this approach to provide initial evidence on whether concept-level semantic representations remain decodable in individuals with chronic post-stroke aphasia.

### Covert feature generation as a clinically relevant semantic task

It is also noteworthy that semantic feature generation is a prominent component of one of the most well studied and commonly used word production treatments for people with aphasia: Semantic Feature Analysis (SFA; Boyle & Coelho, 1995; Coelho et al., 2000). The goal of SFA is to strengthen the connections between *semantic concepts* and their related lexical word forms by guiding a person with aphasia through the production of semantic features related to the target word’s category, physical properties, function, location, and other personal associations. Recent clinical studies demonstrate that both the quantity of generated features (Evans et al., 2021) and their semantic proximity to the target predict naming success in SFA (Cavanaugh et al., 2026), suggesting that semantic feature generation provides a meaningful window into the structure and quality of conceptual representations. Although the present study is not an investigation of SFA per se, the covert feature-generation task offers a lens to better understanding one of the putative key active ingredients of SFA: feature generation. Positioning this task within an fMRI decoding paradigm therefore provides a mechanism for linking theories of semantic representation to clinically relevant behavior.

Building on this conceptual alignment, the present study leverages covert feature generation not as a treatment mechanism, but as a task for directly examining the integrity of conceptual-semantic representations in the brain. Although feature generation is central to SFA and has clear behavioral relevance, no prior work has tested whether concept-level semantic structure, as captured through multivoxel similarity patterns, can be decoded in individuals with aphasia. By applying multivariate neuroimaging methods to a task that engages semantic feature retrieval, the present study addresses this gap and evaluates whether distributed conceptual representations remain identifiable despite stroke-related damage.

First, we used RSA to examine whether neural activation patterns in a theoretically motivated semantic network ROI (Binder et al., 2009) during feature generation aligned with competing semantic models (taxonomic, distributional, and experiential). Second, we applied RSD to test whether concept identity could be accurately recovered from neural similarity structures. We hypothesized that decoding accuracy would be comparable to previous semantic decoding studies using the semantic feature generation paradigm (Anderson et al., 2016; Mitchell et al., 2008) and significantly above chance in people with aphasia, indicating that distributed conceptual representations remain at least partially intact despite stroke-related disruption. Third, we conducted exploratory analyses relating decoding performance in aphasia to standardized language assessments, model-based error-type profiles (Foygel & Dell, 2000), and lesion characteristics. We expected that decoding accuracy would correlate most strongly with semantic measures and lesion load, providing preliminary evidence for links between neural representational integrity and clinically relevant language outcomes. In contrast, given that the semantic models do not contain phonological information and that the scanner task itself does not require accessing phonological word forms, we did not expect that decoding accuracy would not be related to clinical measures of phonological processing.

## Methods

All procedures were approved by the Institutional Review Board at the University of Pittsburgh (Protocol #23040139), and all participants provided written informed consent prior to any study activities. Data collection consisted of clinical assessments of cognition and language followed by structural and functional neuroimaging tasks including semantic feature generation. Clinical assessments were administered by the first author, a licensed speech-language pathologist, in a quiet testing environment before participants underwent any imaging procedures.

### Participants

Six individuals with aphasia participated in this study (age 59–68 years, mean = 62.3 years, SD = 3.4; all right-handed; 2 female; 3 White, 2 Black, 1 American Indian). Time post-onset of aphasia ranged from 90 to 150 months (mean = 114.7, SD = 24.1 months). Lesion volumes ranged from 2.2 to 120.8 cm³ (mean = 52.3 cm³, SD = 51.3), with five ischemic strokes and one hemorrhagic stroke reported. See Figure 1 for lesion overlays from participants with aphasia. Aphasia severity, measured by the Comprehensive Aphasia Test (CAT; Swinburn et al., 2004) Modality Mean T-score, reflected mild to moderate language impairments (range 55.0–61.7, mean = 58.8.7, SD = 2.3). Scores on the Philadelphia Naming Test (PNT; Roach et al., 1996) ranged from 89 to 163 correct out of 174 (mean = 134.8, SD = 30.84). Of note, one PNT item was excluded because its target name is offensive to Alaska Natives. Participants’ lexical-semantic and phonological processing abilities were estimated by fitting their PNT error profiles with the Two-Step Interactive Activation Model (Foygel & Dell, 2000) using a web-based application: https://sites.socsci.uci.edu/~alns/webfit.html. The mean square root s-weight was 0.17 (SD = 0.02), and the mean square root p-weight was 0.16 (SD = 0.02).

**Figure 1.**
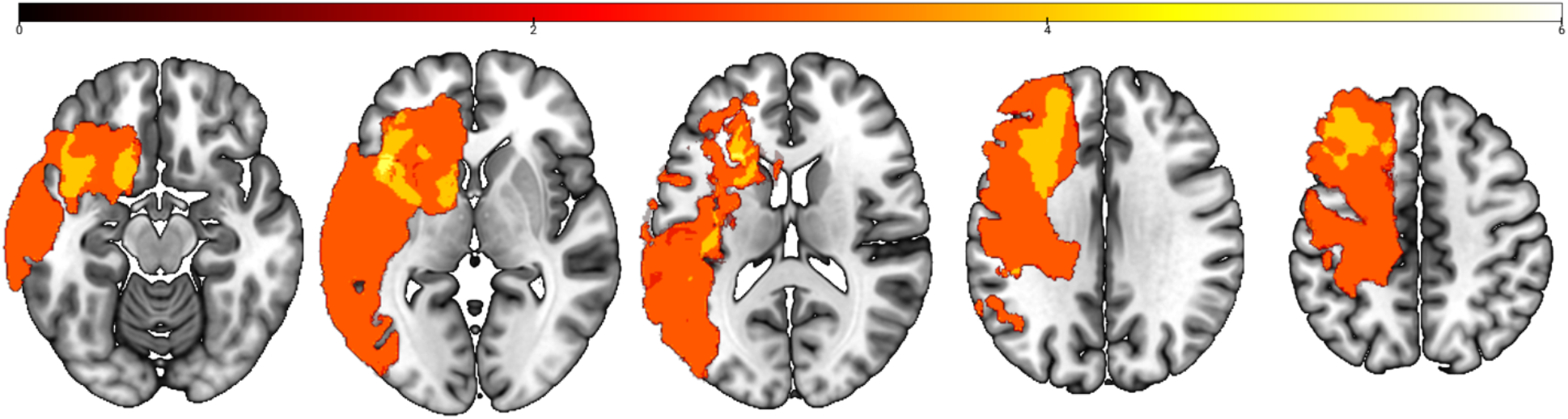
Lesion overlays for participants with aphasia. Darker colors represent lesser overlap of lesions across participants while brighter colors represent greater overlap.

Conceptual semantic processing abilities were measured with the three-picture version of the Pyramids and Palm Trees Test (Howard & Patterson, 1992) (mean proportion correct = 0.94; SD = 0.06; range: 0.82–1.00), Camels and Cactus (Bozeat et al., 2000) (mean proportion correct = 0.85 ; SD = 0.06; range 0.83–0.92), and the Auditory Synonym Judgment subtest from the Psycholinguistic Assessments of Language Processing in Aphasia (PALPA; Kay et al., 1996) (mean proportion correct = 0.85; SD = 0.06; range 0.77–0.90). Phonological processing ability was assessed with the PALPA Nonword Repetition (mean proportion correct = 0.71, SD = 0.17; range: 0.53-0.87), Written Rhyme Judgment (mean proportion correct = 0.90, SD = 0.13; range: 0.63-0.98), and Auditory Rhyme Judgment (mean proportion correct = 0.95, SD = 0.04; range: 0.88-0.98) subtests. Detailed demographic, lesion, and cognitive-linguistic measures are presented in Table 1.

**Table 1.**
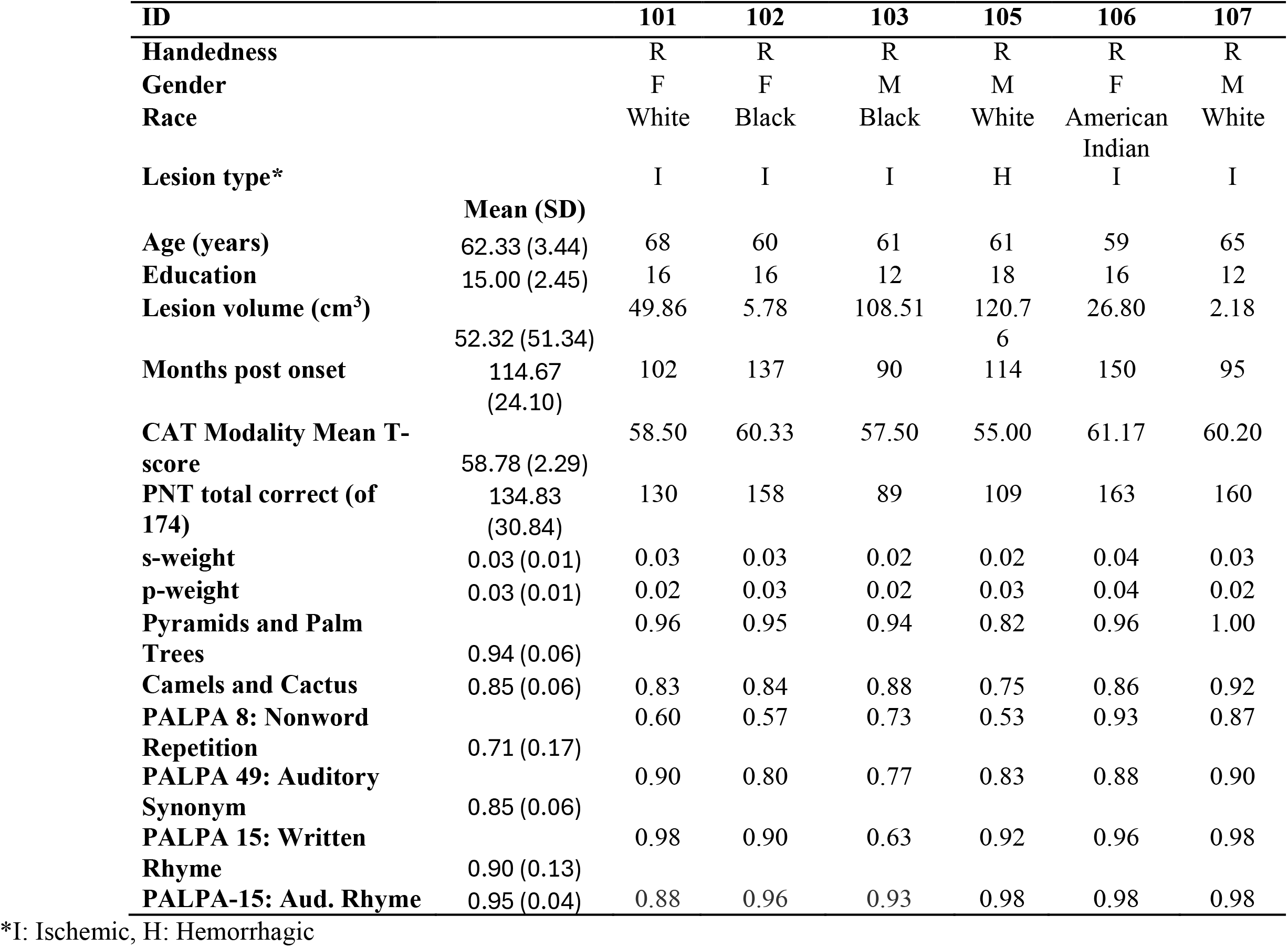
Demographic, lesion information, and cognitive-linguistic data from participants with aphasia.

Eight healthy adults (age 44–72 years, mean = 58.4 years, SD = 11.2; 3 female; 7 right-handed; 6 White, 2 Black) participated as controls. All had normal or corrected-to-normal vision and hearing and completed 12–22 years of education (mean = 15.8 years, SD = 3.6). Cognitive screening with the Saint Louis University Mental Status screener (Morley & Tumosa, 2002) confirmed intact cognitive function (scores 27–30, mean = 28.6, SD = 1.3). Demographic and screening data are provided in Table 2.

**Table 2.** Demographic and cognitive screen results for control participants.

| **ID** | **001** | **002** | **003** | **004** | **005** | **006** | **007** | **008** |
| --- | --- | --- | --- | --- | --- | --- | --- | --- |
| **Handedness** | R | R | R | R | R | R | R | L |
| **Gender** | F | M | M | M | M | F | M | F |
| **Race** | White | White | Black | White | Black | White | White | White |
| **Age** | 45 | 69 | 52 | 53 | 67 | 44 | 72 | 65 |
| **Education** | 16 | 12 | 18 | 18 | 12 | 22 | 12 | 16 |
| **SLUMS** | 30 | 28 | 27 | 30 | 28 | 30 | 29 | 27 |

### Experimental Design and Materials

A dense-sampling design was employed in an effort to obtain reliable item-level estimates of neural activity and enhance sensitivity to within-participant representational patterns (Poldrack, 2017). Each participant was scheduled to complete 12 functional runs of a semantic feature generation task and two high-resolution structural scans for coregistration, normalization, and lesion tracing. However, participants actually completed between 7 and 12 runs, depending on scanner availability, scheduling, and individual comfort. Runs were distributed across 4–5 sessions, with one to three runs completed per session. When possible, sessions were spaced at least one week apart to reduce within-session signal degradation and maximize cross-run reliability (Mazurchuk et al., 2024).

During each functional run, participants viewed 57 image stimuli presented in randomized order. Each trial consisted of an image presentation for four seconds followed by a fixation cross for eight seconds. The images were from six distinct groups: animals (e.g., cheetah), food (e.g., carrot) , humans (e.g., boy), musical instruments (e.g., harp), nature scenes (e.g., tornado), and modes of transportation (e.g., truck). All categories included 10 images, except humans which had 7. Participants were instructed: *“Please silently generate features associated with each image.”* This covert feature-generation paradigm was designed to avoid the use of overt speech while still engaging the core mechanism of SFA, ensuring accessibility for individuals with aphasia. This training protocol follows Mitchell and colleagues (2008) with one exception, the research participants in this study were not encouraged to generate consistent semantic features for each item across runs.

### Semantic feature generation training

Because the primary task required silent semantic feature generation during fMRI, we developed a training protocol to ensure task comprehension in both healthy controls and participants with aphasia. Participants were first introduced to the distinction between strong features, defined as specific, diagnostic attributes likely to aid retrieval (e.g., target: *rose*, feature: *flower*), and weak features, defined as overly general or idiosyncratic responses (e.g., target: *rose*, feature: *nice*). During familiarization, each item was presented in a slideshow with its written and spoken label, and participants were asked to generate at least one semantic feature per item. Weak responses were corrected with feedback to reinforce the distinction between strong and weak features.

To simulate in-scanner demands, participants then completed a PsychoPy-based (Peirce, 2007) practice block that matched the timing of the fMRI task (4 second image presentation, 8 second fixation). Participants pressed a button after silently generating a feature for each image, either during the image presentation or the fixation interval. Performance was defined as the proportion of trials with a button press, indexing task compliance rather than feature accuracy because feature generation occurred covertly. All participants responded on at least 90% of trials on their first attempt, indicating reliable task engagement, although several individuals with aphasia requested additional practice, which occurred prior to participation in the fMRI experiment.

### fMRI data acquisition

All neuroimaging data were collected at the Carnegie Mellon University-University of Pittsburgh Brain Imaging Data Generation & Education (BRIDGE) Center with a 3T Siemens Prisma scanner equipped with a 64-channel Head/Neck coil. The first scanning session began with a high-resolution T1-weighted anatomical image acquired using a magnetization-prepared rapid acquisition gradient echo (MPRAGE) sequence (TR = 2400 ms; TE = 2.22 ms; flip angle = 8°; voxel size = 1.0 × 1.0 × 1.0 mm; 256 × 256 matrix; 176 slices; 1 mm slice thickness), followed by a T2-weighted FLAIR scan for anatomical validation and lesion identification (TR = 5000 ms; TE = 395 ms; flip angle = 120°; voxel size = 1.0 × 1.0 × 1.0 mm). A posterior–anterior phase-encoded EPI reference acquisition was also collected for susceptibility distortion correction (TR = 2000 ms; TE = 30 ms; voxel size = 2.0 × 2.0 × 2.0 mm). Susceptibility-induced distortions were estimated from EPI references with opposing phase-encoding directions using FSL TOPUP. Following the structural sequences and field maps, the feature generation functional tasks were completed. Functional images were collected using a multiband gradient-echo echo-planar imaging (EPI) sequence (TR = 1000 ms; TE = 30 ms; flip angle = 55°; multiband factor = 4; 66 axial slices; voxel size = 2.0 × 2.0 × 2.0 mm). Slice acquisition was performed using interleaved ordering. Each functional run lasted 11.4 minutes, corresponding to 684 volumes,.

### fMRI data preprocessing and first-level analysis

Preprocessing was conducted with fMRIPrep 20.2.0 (Esteban et al., 2019), which implements best practices in fMRI preprocessing using Nipype (Gorgolewski et al., 2011). For anatomical images, T1-weighted volumes were corrected for intensity non-uniformity, skull-stripped, and nonlinearly registered to the ICBM152 Nonlinear Asymmetrical template (Fonov et al., 2009) using ANTs (Avants et al., 2008). Brain tissue was segmented into gray matter, white matter, and cerebrospinal fluid (Zhang et al., 2001). For participants with aphasia, lesions were manually traced in ITK-SNAP and binary lesion masks were incorporated into spatial normalization to improve alignment across individuals.

Functional data were slice-time corrected (AFNI; Cox, 1996), motion corrected (FSL MCFLIRT; Jenkinson et al., 2002), and distortion corrected using the TOPUP method (Andersson et al., 2003). Participants’ functional runs were co-registered to their T1-weighted image using boundary-based registration with 12 degrees of freedom (Greve & Fischl, 2009). Anatomical noise regressors were estimated with aCompCor (Behzadi et al., 2007), and six principal components were retained from white matter and CSF masks. Framewise displacement was computed for each run (Power et al., 2014).

First-level analyses were implemented in Nilearn’s FirstLevelModel (Abraham et al., 2014). The design matrix for each run included a separate regressor for every individual stimulus (i.e., one per stimulus picture). Each stimulus was modeled as a 12s boxcar function beginning at image onset and spanning the entire trial, including the 4s picture presentation and the subsequent 8-s fixation interval, spanning the full semantic feature generation epoch. Stimulus regressors were convolved with the canonical SPM hemodynamic response function. Nuisance regressors were derived from the fMRIPrep confound outputs and included the six rigid-body motion parameters, their temporal derivatives, six anatomical CompCor components, and outlier volumes exceeding 0.7 mm framewise displacement. Models also included cosine drift terms (high-pass cutoff = 128 s). A 6 mm FWHM Gaussian smoothing kernel was applied during model estimation. For each run, item-level beta maps were computed and saved separately, providing run-specific, stimulus-level neural activation patterns for subsequent representational similarity and decoding analyses.

### Region of interest

The semantic network ROI (Figure 2) corresponded to the top 1% most consistently activated voxels across 120 prior fMRI studies of semantic language processing, as originally compiled by Binder et al. (2009). This data-driven ROI captures the distributed set of regions most reliably engaged in conceptual processing, including portions of the inferior frontal gyrus, middle and superior temporal gyri, angular gyrus, and ventral temporal cortex. The ROI was binarized and resampled to each participant’s native functional space prior to voxel extraction, and lesioned voxels were excluded for participants with aphasia. This ROI served as the primary focus of all RSA analyses.

**Figure 2.**
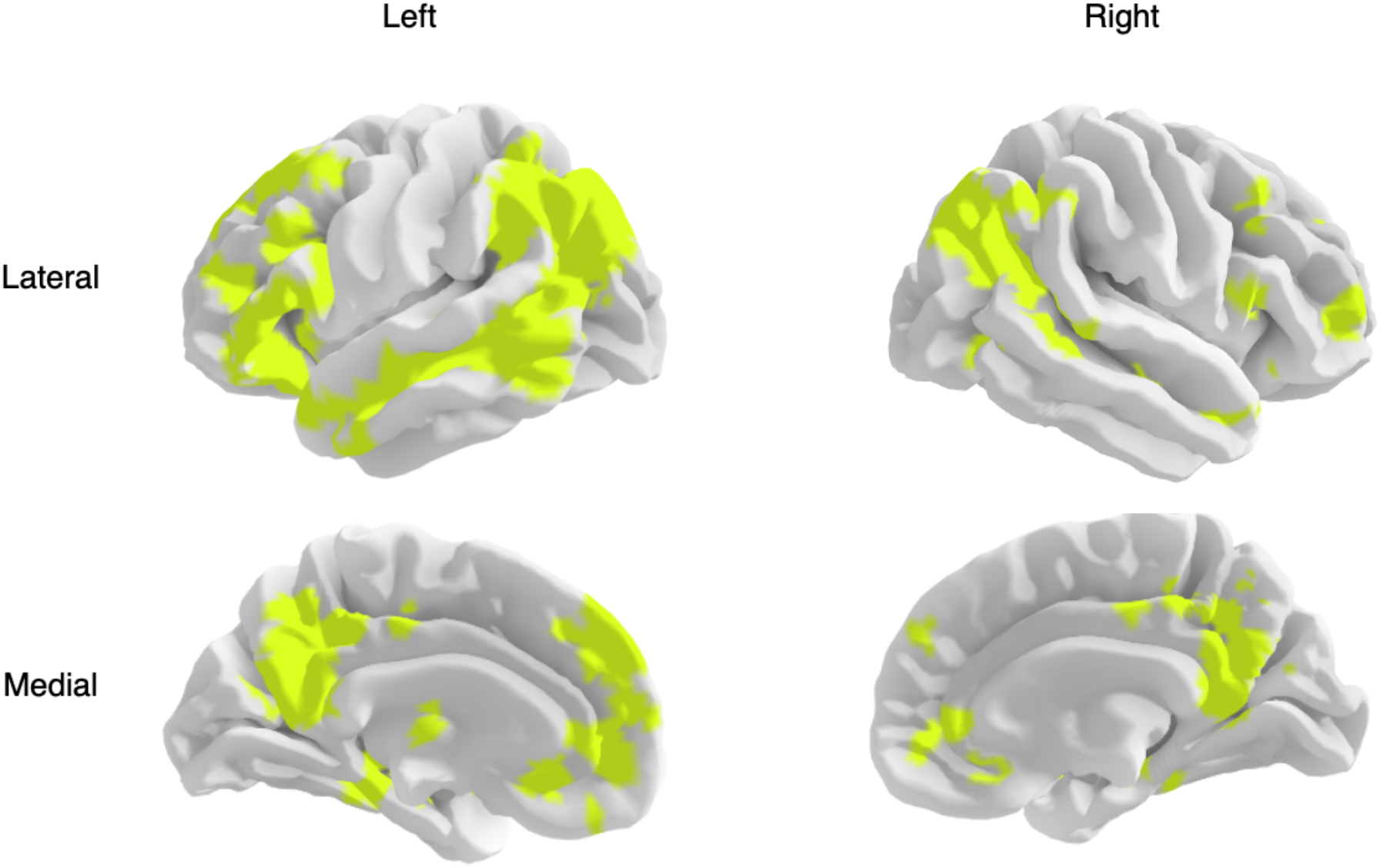
Semantic Network Region of Interest included in the RSA analysis Temporal signal-to-noise ratio (tSNR)

Voxel-level signal quality was evaluated using temporal signal-to-noise ratio (tSNR), defined as the mean signal over time divided by its standard deviation (Welvaert & Rosseel, 2013). For each participant, tSNR was computed from preprocessed, unsmoothed 4D BOLD time series prior to model estimation. Voxels falling below the 30th percentile of the participant’s tSNR distribution were excluded. The remaining voxels comprised a participant-specific tSNR mask, representing those voxels with the highest temporal signal reliability. This thresholding approach is consistent with prior work showing that moderate tSNR filtering improves multivariate reliability while preserving critical information (Gardumi et al., 2016).

### Neural Representational Dissimilarity Matrices (RDMs)

For each participant, voxel masks were defined as the intersection of the semantic network ROI, the participant-specific tSNR mask, and (for participants with aphasia) manually traced lesion masks. Within these masks, beta maps were averaged across all available runs for each of the 57 stimulus items. Item-wise activation patterns were then vectorized, and dissimilarities were computed as one minus the Pearson correlation across voxels, yielding a 57 × 57 symmetric neural RDM for each participant. Group-level RDMs were generated by averaging participant-level RDMs within each group.

### Split-Half Reliability (Noise Ceiling)

To estimate the internal consistency of neural RDMs, we computed participant-level split-half reliability across runs. For each iteration, available runs were randomly shuffled and divided into two non-overlapping subsets. Within each subset, item-level beta maps were averaged across runs for each item, and a neural RDM was constructed separately for each subset. Reliability was quantified as the Pearson correlation between the upper-triangular elements of the two RDMs, excluding the diagonal. This procedure was repeated across 100 random run partitions, and the mean correlation across splits served as the reliability estimate for each participant. These estimates provided participant-level noise ceilings for interpreting model–neural RSA correlations.

### Semantic Model RDMs

We constructed RDMs based on four semantic models spanning three theoretical domains: taxonomic, distributional, and experiential (Figure 3). The taxonomic model was based on WordNet (Fellbaum, 2010), a lexical database organizing English words into hierarchical synsets. Pairwise similarity was computed using the Wu & Palmer metric (Wu & Palmer, 1994), which considers the depth of two concepts and their most specific common ancestor. Dissimilarity for the WordNet model was defined as 1 – Wu & Palmer similarity metric.

**Figure 3.**
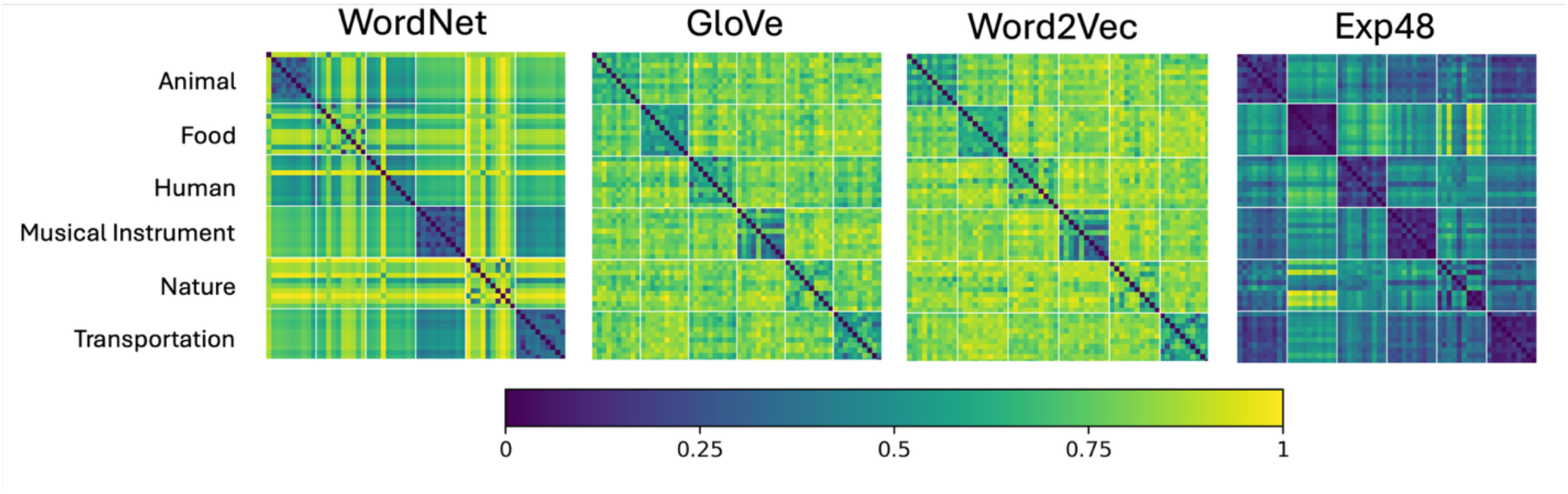
Heatmaps of pairwise dissimilarities for each tested semantic model. Each column and row represents one of the 57 concepts used in the study semantic distance to another concept. Word pair relationships with darker colors represent model estimated with lesser disimilairty in meaning compared to those with brighter hues. For visual clarity, words within each category are grouped together.

Two distributional models captured semantic structure from word co-occurrence statistics. GloVe (Pennington et al., 2014) embeddings were derived from global co-occurrence ratios whereas Word2Vec (Mikolov et al., 2013) used a Skip-Gram architecture to predict local context words. Both were implemented with pre-trained 300-dimensional embeddings, and dissimilarities were defined as 1 minus the Pearson correlation between vectors.

One experiential model (Exp48), which represents concepts based on human ratings of perceptual, motor, affective, and contextual features (Binder et al., 2016) was used in the study. Exp48 captures 48 dimensions spanning sensory, motor, spatial, temporal, and affective-cognitive experiences. Dissimilarity was also defined for the Exp48 model as 1 minus the Pearson correlation between vectors.

To control for potential low-level visual similarity, we also constructed a visual similarity RDM. Stimulus images were passed through a pretrained ResNet50 convolutional neural network (He et al., 2015), and 2048-dimensional embeddings capturing high-level perceptual features such as texture, contour, and shape were extracted from the penultimate average pooling layer. The resulting visual embeddings were then L2-normalized (i.e., scaled to unit length) to standardize vector magnitude. Pairwise dissimilarities were then computed using cosine distance.

### Representational Similarity Analysis

To account for low-level perceptual similarity, neural dissimilarities were regressed on visual dissimilarities using a simple linear model, and the residuals were retained as the visually corrected neural RDM. This correction was applied prior to both raw and partial RSA. In the raw RSA, each semantic model RDM (WordNet, Word2Vec, GloVe, and Exp48) was correlated with the residualized neural RDM using Spearman correlation. In the partial RSA, the unique contribution of each semantic model was estimated while controlling for variance explained by the remaining models. For example, the unique contribution of Exp48 was quantified by computing the partial Spearman correlation between the neural RDM and the Exp48 RDM while controlling for WordNet, Word2Vec, and GloVe. Together, these analyses allowed us to evaluate both overall model–brain alignment and model-specific contributions to neural representational structure. Group-level significance of participant-level RSA values was assessed using Wilcoxon signed-rank tests against zero. To control for multiple comparisons, p values were adjusted using false discovery rate correction (q = 0.05) separately within each group and RSA type across the four semantic models.

## Representational Similarity Decoding

### Voxel selection and neural RDM construction

For each participant, we selected the 500 most temporally stable voxels from the full gray matter volume. Gray matter masks were derived from probabilistic tissue segmentations generated by fMRIPrep and binarized to retain voxels with nonzero gray matter probability. For participants with aphasia, voxels overlapping manually traced lesion masks were excluded so that only viable tissue contributed to decoding analyses. Temporal stability was quantified from cross-run correlations of trial-level beta estimates. For each voxel, we extracted a trial × run activation matrix, computed a similarity matrix across runs, and averaged the lower triangle to obtain a voxel-wise stability score. The 500 voxels with the highest average stability were retained for subsequent decoding. Within this selected voxel set, beta estimates were averaged across all available runs for each item, yielding one activation vector per stimulus. Neural RDMs for decoding were then computed as one minus the Pearson correlation between all item vectors.

### Semantic model selection and visual control

Exp48 was selected as the primary semantic model for decoding because it showed the strongest correspondence with neural representational structure in the preceding RSA analyses across both groups. As in the RSA analyses, visual similarity was controlled for with ResNet-50 and was regressed from neural RDMs prior to semantic decoding.

### Leave-two-out decoding procedure

Decoding was implemented using a leave-two-out procedure (Anderson et al., 2016; Figure 4). On each fold, two items were held out as test stimuli, and reduced neural and semantic RDMs were computed after removing information corresponding to the held-out pair. For each held-out item, a neural similarity vector was extracted and compared with the candidate semantic vectors. A decoding trial was scored as correct when the sum of correlations under the correct item assignment exceeded that under the swapped assignment. Accuracy was calculated as the proportion of correct trials across all 1,596 possible item pairs.

**Figure 4.**
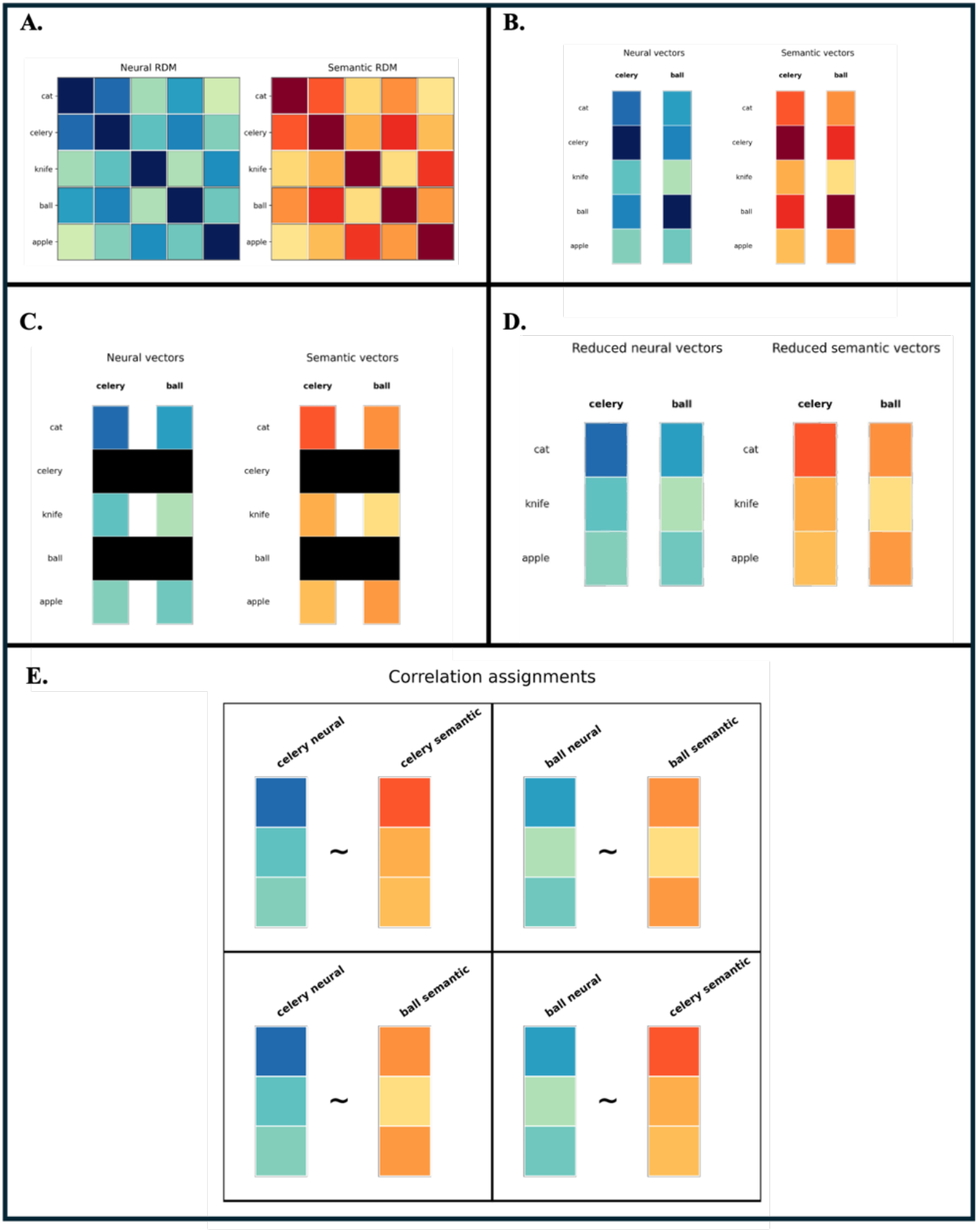
Schematic of the leave-two-out representational similarity decoding procedure. **A:** Representational dissimilarity matrices are estimated from the neural data (cool hues) and the semantic model (warm hues). **B:** Two words are selected for decoding, and their corresponding vectors are extracted from each RDM. **C:** The self-correlations and mutual correlations between the two held-out words are identified. **D:** These elements are removed to prevent information leakage, yielding reduced vectors for decoding. **E:** Neural and semantic vectors are then compared under the two possible assignments. Decoding is considered successful when the sum of the two true correlations exceeds the sum of the two false correlations.

### Permutation-based null distributions

Statistical significance was assessed using participant-specific permutation tests. For each participant, 1,000 null semantic RDMs were generated by randomly permuting item labels while preserving the structure of the neural RDM. Each permuted RDM was submitted to the same leave-two-out decoding procedure, yielding a null distribution of accuracies. *P*-values were calculated as the proportion of null accuracies greater than or equal to the observed accuracy.

### Grid search for joint model contributions

Although Exp48 showed the strongest correspondence with neural representational structure in the RSA analyses (see below), taxonomic and distributional models may nonetheless capture complementary aspects of semantic organization. To test this possibility, we conducted a grid search combining the experiential (Exp48), taxonomic (WordNet), and top performing distributional (Word2Vec) RDMs. Model weights were varied in 0.05 increments across all possible mixtures, constrained to sum to one. For each weight configuration, a joint semantic RDM was computed as a weighted linear combination of the three z-scored model RDMs. Each joint RDM was then entered into the leave-two-out decoding procedure, yielding an accuracy surface across the weight space. The optimal model was defined as the weight combination that maximized mean decoding accuracy across participants.

## Results

### Voxel retention and data quality

To ensure high-quality voxel-level data for RSA, we computed tSNR maps for each participant from preprocessed, unsmoothed BOLD time series. Voxel inclusion was restricted to the top 70% of tSNR values within each participant’s brain mask, and only voxels overlapping the functionally defined semantic network ROI were retained for RSA analyses. For participants with aphasia, voxels within manually traced lesion masks were also excluded to prevent spurious correlations.

Participants with aphasia retained an average of 8,679 voxels (SD = 1,123), ranging from 7,219 to 9,750. Healthy controls retained an average of 9,568 voxels (SD = 406), ranging from 8,770 to 10,033. This group difference was not significant, t(5.99) = 1.85, p = .11. The lower voxel retention in participants with aphasia likely reflects expected signal degradation and anatomical disruption associated with stroke. Nonetheless, all included participants exceeded the predefined threshold for multivoxel analysis (i.e., retention of voxels above the 30th percentile of the participant-specific tSNR distribution).

The number of completed functional runs did not differ significantly between groups, t(9.88) = 0.66, p = .53. Participants with aphasia completed an average of 9.8 runs (SD = 2.1, range = 7–12), whereas controls completed an average of 10.5 runs (SD = 1.8, range = 7–12).

### Representational similarity analysis within the semantic network ROI: Participants with aphasia

#### Raw correlations

At the participant level, participants with aphasia (Figure 5) showed the strongest raw RSA correspondence with Exp48. Raw Exp48 RSA correlation values were positive in all six participants (mean = 0.13, range = 0.02 to 0.29), and participant-level raw RSA values were significantly greater than zero across participants before correction for multiple comparisons (Wilcoxon signed-rank p = 0.03). By comparison, raw RSA effects for the other models were weaker and more variable. Participants with aphasia showed weaker raw RSA correspondence for Word2Vec (mean = 0.08, range = -0.02 to 0.28) and GloVe (mean = 0.06, range = -0.05 to 0.23), and WordNet showed essentially no correspondence overall (mean = 0.01, range = -0.07 to 0.08). Raw RSA values for Word2Vec, GloVe, and WordNet were not significantly greater than zero across participants (all Wilcoxon signed-rank ps ≥ 0.22). After FDR correction across the four model tests within the aphasia raw RSA analysis, no model remained significant.

**Figure 5.**
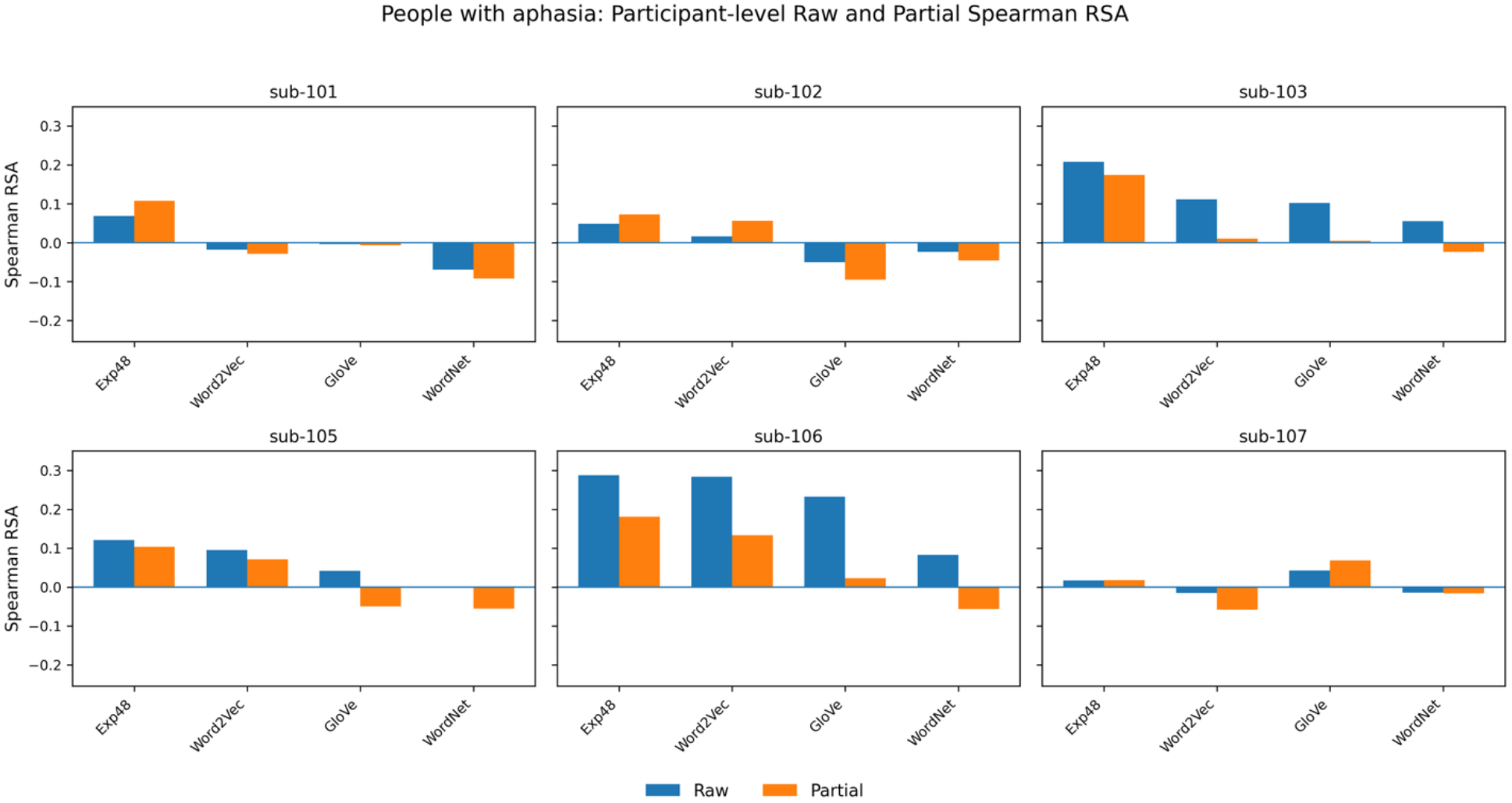
Participant-level semantic network RSA results (Participants with aphasia). Blue (left columns) represent raw correlations and orange (right) represent partial correlations.

#### Partial correlations

The same overall pattern was observed for partial RSA. After controlling for the other semantic models, Exp48 remained positive in all six participants (mean = 0.11, range = 0.02 to 0.18), and participant-level partial RSA values were significantly greater than zero across participants before correction for multiple comparisons (Wilcoxon signed-rank p = 0.03). In contrast, partial RSA effects for the other models were smaller and did not show comparable evidence of positive unique correspondence. Participants with aphasia showed little to no partial RSA correspondence for Word2Vec (mean = 0.03, range = -0.06 to 0.13) and GloVe (mean = - 0.01, range = -0.09 to 0.07), whereas WordNet partial RSA values were negative overall (mean = -0.05, range = -0.09 to -0.02). Partial RSA values for Word2Vec and GloVe were not significantly greater than zero across participants (Wilcoxon signed-rank ps ≥ 0.44), whereas WordNet partial RSA values were significantly less than zero before correction (Wilcoxon signed-rank p = 0.03). After FDR correction across the four model tests within the aphasia partial RSA analysis, no model remained significant.

### Representational similarity analysis within the semantic network ROI: Healthy controls

#### Raw correlations

At the participant level, healthy controls (Figure 6) also showed the strongest raw RSA correspondence with Exp48. Raw Exp48 RSA values were positive in all eight control participants (mean = 0.13, range = 0.03 to 0.26), and participant-level raw RSA values were significantly greater than zero across participants (Wilcoxon signed-rank p = 0.01). By comparison, raw RSA effects for the other models were weaker and more variable. Controls showed weaker raw RSA correspondence for Word2Vec (mean = 0.08, range = -0.01 to 0.22) and GloVe (mean = 0.05, range = -0.04 to 0.16), whereas WordNet showed little to no correspondence (mean = -0.02, range = -0.11 to 0.06). Raw RSA values for Word2Vec, GloVe, and WordNet were not significantly greater than zero across participants (all Wilcoxon signed-rank ps ≥ 0.08). After FDR correction across the four model tests within the control raw RSA analysis, only Exp48 remained significant.

**Figure 6.**
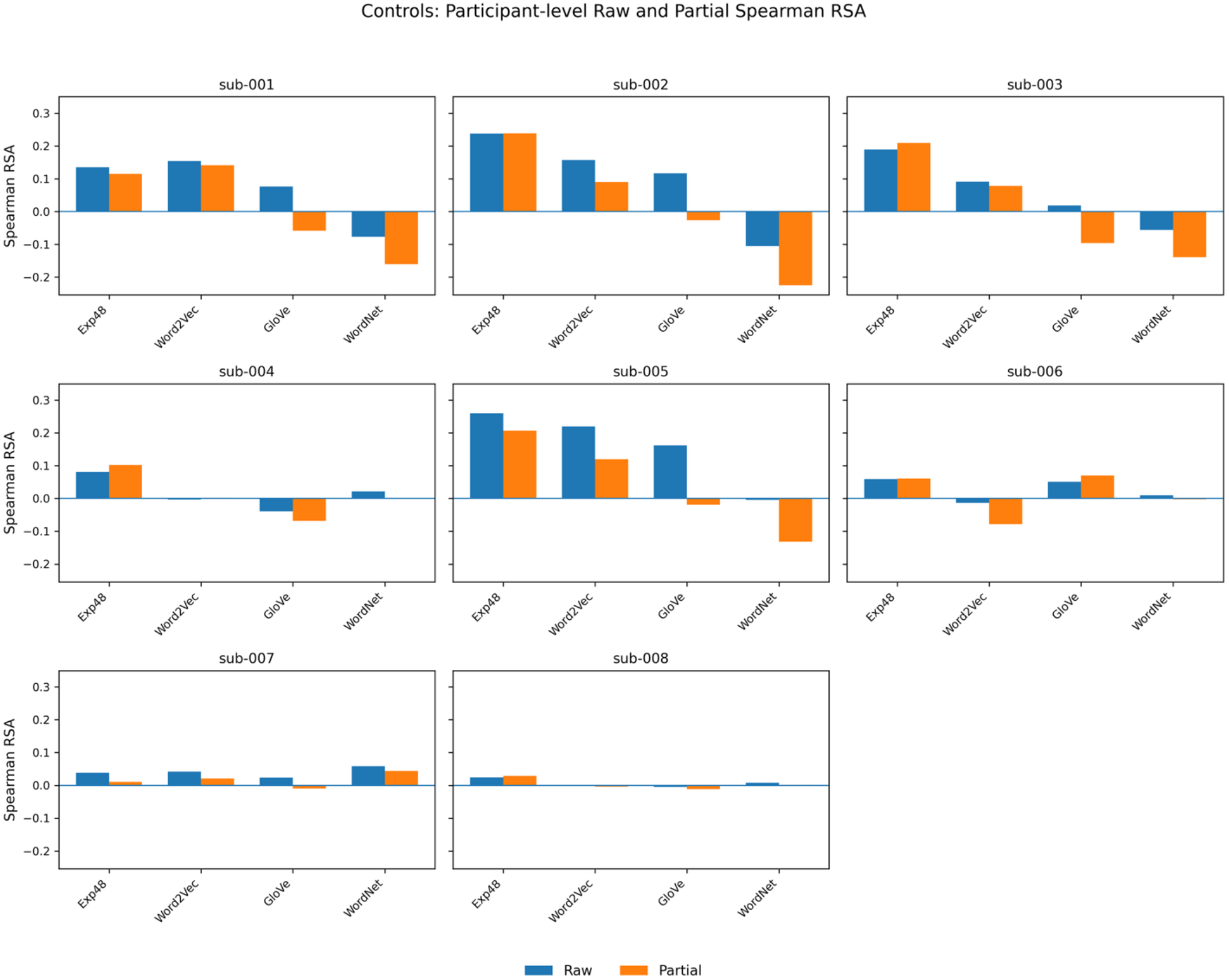
Group-level Representational similarity results (Control group). Blue (left columns) represent raw correlations and orange (right) represent partial correlations.

#### Partial correlations

The same overall pattern was observed for partial RSA. After controlling for the other semantic models, Exp48 remained positive in all eight control participants (mean = 0.12, range = 0.01 to 0.24), and participant-level partial RSA values were significantly greater than zero across participants (Wilcoxon signed-rank p = 0.01). In contrast, partial RSA effects for the other models were smaller and did not show comparable evidence of positive unique correspondence. Controls showed weaker partial RSA correspondence for Word2Vec (mean = 0.05, range = -0.08 to 0.14) and GloVe (mean = -0.03, range = -0.10 to 0.07), whereas WordNet partial RSA values were negative overall (mean = -0.08, range = -0.22 to 0.04). Partial RSA values for Word2Vec, GloVe, and WordNet were not significantly greater than zero across participants (all Wilcoxon signed-rank ps ≥ 0.11). After FDR correction across the four model tests within the control partial RSA analysis, only Exp48 remained significant.

### Representational similarity decoding results

#### Representational Similarity Decoding with Experiential Semantic Features (Exp48)

We next evaluated decoding accuracy using the Exp48 semantic model, which was selected because it demonstrated the strongest unique correspondence with neural representational geometry in the RSA analyses. As with the RSA analyses reported above, we controlled for visual similarity by residualizing the neural RDM using the visual model prior to conducting the decoding analyses. after controlling for variance explained by competing models. In the aphasia group, mean decoding accuracy was 71.03% (range: 61.78–78.95%), with three of six participants exceeding 75% decoder accuracy. Detailed individual accuracies are provided in Table 3.

**Table 3:** Participant-level RSD decoding accuracies and permutation p-values (people with aphasia).

| Participant | Accuracy | Permutation<br>p-value |
| --- | --- | --- |
| sub-101 | 61.78 | 0.049 |
| sub-102 | 64.41 | 0.014 |
| sub-103 | 78.13 | 0.001 |
| sub-105 | 73.37 | 0.001 |
| sub-106 | 78.95 | 0.001 |
| sub-107 | 69.55 | 0.002 |
| Mean | 71.03 |  |
| SD | 7.07 |  |

In the control group, mean decoding accuracy based on Exp48 was 69.43% (range: 49.5–84.84%). Four of the eight participants exceeded 75% decoding accuracy. However, participants 006, 007, and 008 performed near chance, substantially lowering the overall group mean. Individual accuracies and permutation p values are provided in Table 4.

**Table 4.** Participant-level RSD decoding accuracies and permutation p-values (controls).

| Participant | Accuracy | Permutation<br>p-value |
| --- | --- | --- |
| sub-001 | 78.07 | 0.001 |
| sub-002 | 80.26 | 0.001 |
| sub-003 | 84.84 | 0.001 |
| sub-004 | 68.92 | 0.003 |
| sub-005 | 84.59 | 0.001 |
| sub-006 | 49.5 | 0.525 |
| sub-007 | 52.94 | 0.340 |
| sub-008 | 50.44 | 0.509 |
| Mean: | 68.70 |  |
| SD: | 15.51 |  |

#### Grid search of model contributions

A grid search was conducted to determine if a linear combination of semantic feature model RDMs (rather than an RDM based on one model) would optimize decoding accuracy. The grid search indicated that decoding performance in both groups was optimized when relying primarily on experiential features, with smaller and more variable contributions from distributional and taxonomic dimensions across participants. In participants with aphasia (Figure 7), the optimal mixtures placed the greatest average weight on Exp48 (mean w = 0.67, SD = 0.44), with smaller contributions from Word2Vec (mean w = 0.21, SD = 0.33) and WordNet (mean w = 0.12, SD = 0.21). Mean decoding accuracy for the optimized mixtures (72.5%, SD = 8.6) was overall similar to the mean accuracy obtained using Exp48 alone.

**Figure 7:**
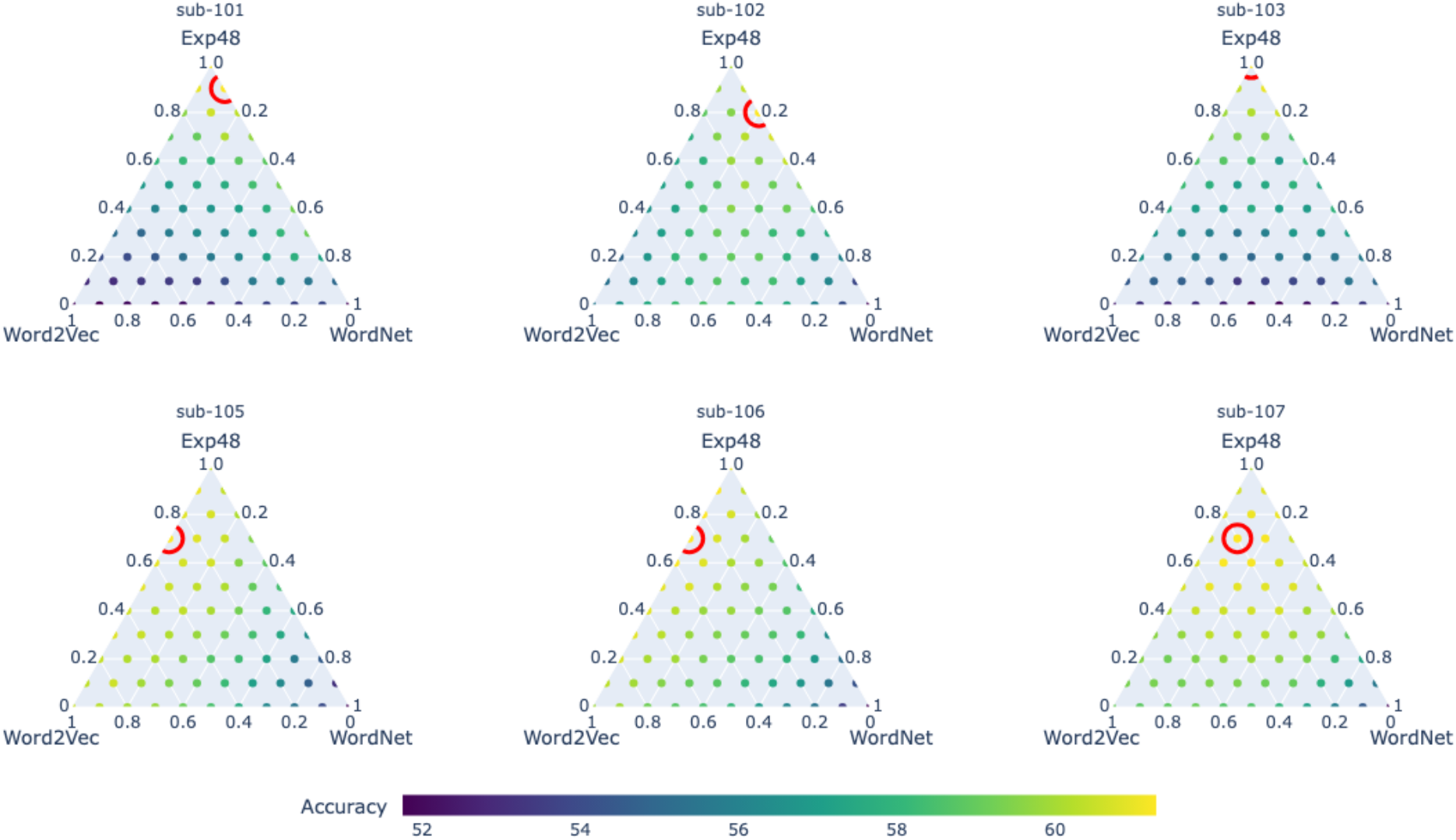
Ternary plot of grid search decoding accuracies for participants with aphasia. Increasing values on the left diagonal from bottom to top represent increases in decoder accuracy as model weight for Exp48 increase. Values increase from top to bottom on the right diagonal of the triangle for model weights for Wordnet. The base of the triangle represents model weights for Word2Vec and increase from right to left. The plot points depict accuracy from low to high with cold to hot colors. For each prediction space, the optimal model weights from the three models are circled in red.

A similar pattern emerged in healthy controls (Figure 8). The optimal weight combinations assigned the greatest average weight to the experiential model (Exp48; mean w = 0.69, SD = 0.35), followed by Word2Vec (mean w = 0.25, SD = 0.38) and minimal contribution from WordNet (mean w = 0.06, SD = 0.12). The corresponding mean decoding accuracy for the optimized mixtures (69.4%, SD = 15.9) was not significantly different from the mean accuracy obtained using Exp48 alone. Taken together, these findings indicate that experiential information provided the dominant basis for neural decodability in participants with aphasia and healthy controls. Although some individuals showed optimal performance with mixed model weights, there was no evidence that adding taxonomic or distributional information improved decoding accuracy at the group level beyond Exp48 alone.

**Figure 8:**
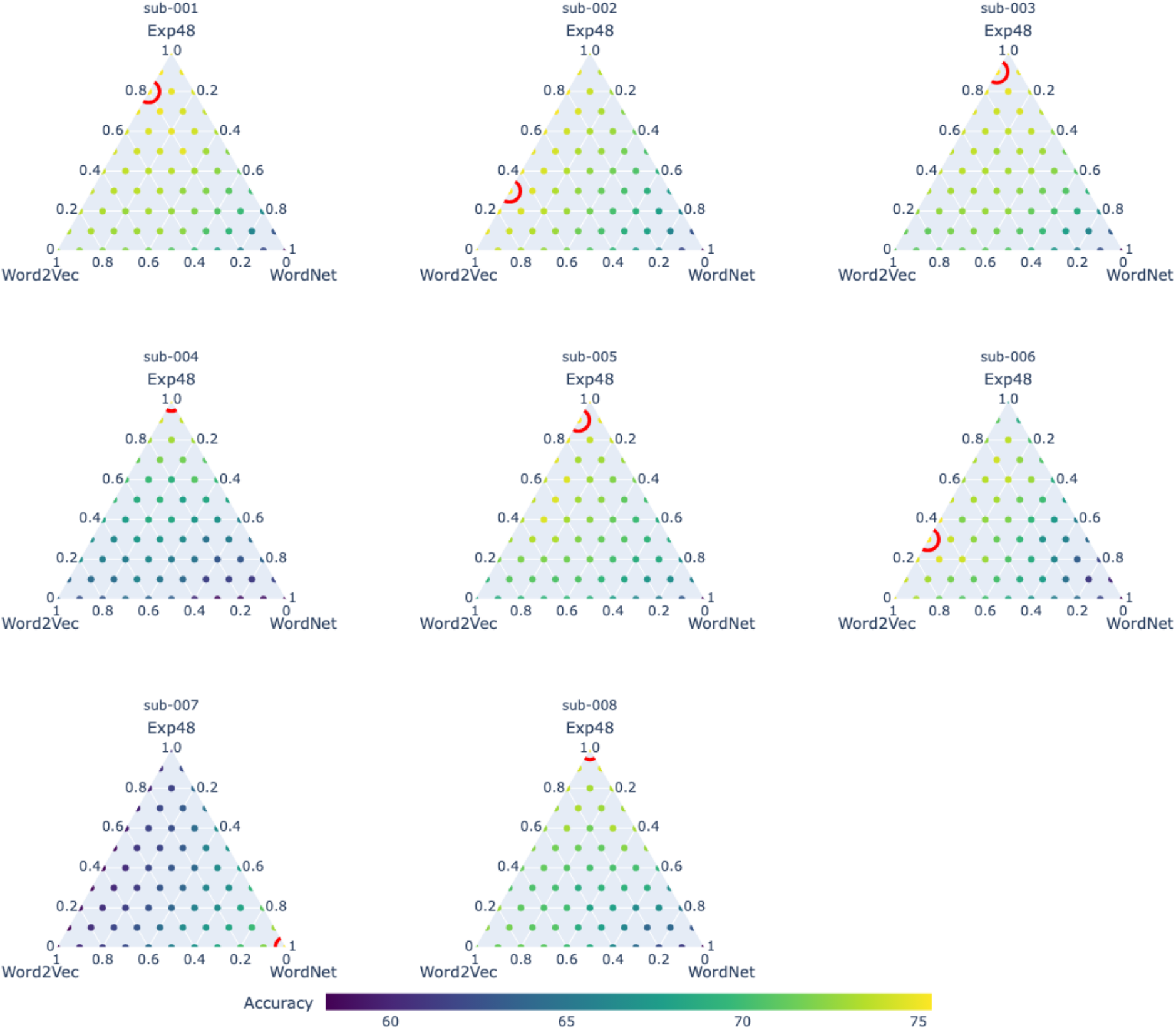
Ternary plot of grid search decoding accuracies for control participants. Increasing values on the left diagonal from bottom to top represent increases in decoder accuracy as model weight for Exp48 increase. Values increase from top to bottom on the right diagonal of the triangle for model weights for Wordnet. The base of the triangle represents model weights for Word2Vec and increase from right to left. The plot points depict accuracy from low to high with cold to hot colors. For each prediction space, the optimal model weights from the three models are circled in red.

#### Anatomical locations of 500 stable voxels

To characterize the anatomical distribution of voxels contributing to representational similarity decoding (RSD), each participant’s 500 most temporally stable voxels within the semantic network mask were mapped onto the Automated Anatomical Labeling (AAL) atlas. Group-level voxel overlap maps are shown in Figure 9, and complete AAL distributions are provided in Supplementary Table 1 and 2.

**Figure 9:**
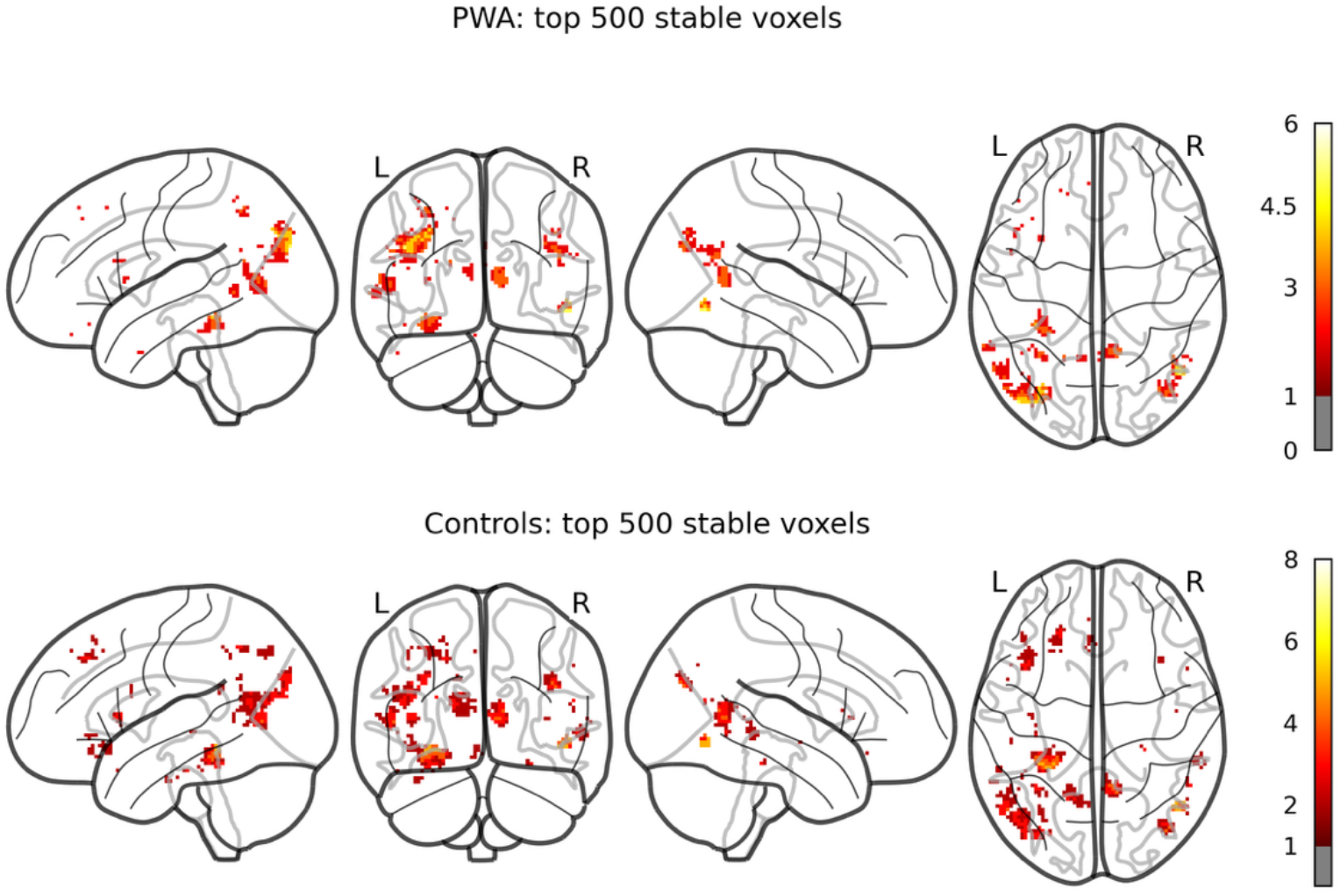
Anatomical distribution of voxels included in representational similarity decoding analyses. Maximum intensity projection maps displaying the spatial overlap of each participant’s 500 most temporally stable voxels within the semantic network mask. Separate overlap maps are shown for participants with aphasia (top) and healthy controls (bottom). Voxel intensity reflects the number of participants within each group for whom that voxel was selected.

In participants with aphasia, the largest voxel counts were observed in the left middle occipital gyrus, left middle temporal gyrus, left precuneus, right angular gyrus, and left fusiform gyrus. Additional voxels were distributed across bilateral temporal, occipital, and inferior parietal regions.

In healthy control participants, the largest voxel counts were observed in the left middle temporal gyrus, left fusiform gyrus, left middle occipital gyrus, left precuneus, and left parahippocampal gyrus. Additional voxels were distributed across bilateral temporal, occipital, and inferior parietal regions.

#### Behavioral and clinical correlates of decoder accuracy

To examine whether individual differences in language ability were associated with decoding performance, we tested relationships between decoder accuracy and a set of clinical and psycholinguistic measures in participants with aphasia. These included aphasia severity (CAT Modality Mean T-score), lexical-semantic processing (s-weight, Pyramids and Palm Trees, Camels and Cactus), and phonological processing (p-weight, PALPA nonword repetition, PALPA rhyme judgments). None of the semantic measures, phonological measures, aphasia severity scores, or lesion volume were significantly associated with decoder accuracy (all *p-* values > .15).

#### Data quality correlates of decoder accuracy

We examined whether decoder accuracy was associated with two indices of fMRI data quality: the number of functional runs acquired and the split-half reliability of the BOLD signal, a measure of within-participant neural reliability. Because these variables were not specific to diagnostic group, this analysis combined all participants (*n* = 14; 8 controls, 6 participants with aphasia). Spearman correlations revealed positive associations between decoder accuracy and both data quality measures. Decoder accuracy was strongly correlated with the noise ceiling (ρ = 0.84, *p* < .001), indicating that participants with more reliable neural similarity structure achieved higher decoding performance. Decoder accuracy was also positively correlated with the number of functional runs completed (ρ = 0.61, *p* = .021).

## Discussion

The present study examined whether concept-level semantic information is decodable using task-based fMRI data in individuals with chronic post-stroke aphasia and how different theoretical models of semantic representation correspond to neural activity patterns during semantic feature generation. Using dense-sampling fMRI combined with RSA and RSD, we evaluated both the structure of semantic information in distributed neural activation patterns and whether that structure supported item-level identification after stroke-induced aphasia. Three primary findings emerged. First, for individuals with aphasia and healthy controls, experiential semantic features (Exp48) showed the strongest correspondence with neural representational geometry, outperforming taxonomic and distributional models in both raw and partial RSA. Second, concept identity could be decoded in both groups using experiential semantic similarity, with decoding accuracy above chance and broadly comparable across both groups. Third, in participants with aphasia and healthy adults, the 500 most stable voxels supporting decoding were distributed across temporal, occipital, and inferior parietal regions, with partially overlapping anatomical distributions across groups. Together, these findings demonstrate that conceptual-semantic representational structure remains observable and can support concept decoding in chronic aphasia and that experiential information provides the strongest account of the underlying neural representations.

This study provides initial evidence that word-level concepts are decodable in people with aphasia. Furthermore, the RSA results provide insight into the type of semantic information that most consistently accounts for neural similarity patterns during feature generation. At the participant level, Exp48 showed the most consistent correspondence with neural representational geometry in participants with aphasia: raw and partial RSA values were positive in all six individuals, whereas taxonomic and distributional models showed weaker and more variable effects. Although Exp48 effects in the aphasia group were significantly greater than zero before correction for multiple comparisons, they did not survive FDR correction and therefore should be interpreted cautiously. A similar but statistically stronger pattern was observed in healthy adults, where Exp48 was positive in all eight participants and remained significant after FDR correction in both raw and partial RSA. Together, these findings suggest that, compared with other models tested, experiential semantic structure provides the most reliable account of neural similarity patterns during covert feature generation, including in chronic post-stroke aphasia. By contrast, distributional models contributed weaker and less consistent correspondence, whereas WordNet showed little raw correspondence and negative partial RSA values once shared variance with other models was controlled. These findings converge with prior RSA work showing prominent experiential representations in heteromodal regions of the semantic network (Fernandino et al., 2022; Tong et al., 2022) and extend this evidence by showing that experiential semantic structure is also detectable at the participant level in a clinical aphasia sample.

A central question motivating this work was whether stroke-induced damage disrupts the neural representational structure of conceptual knowledge to the extent that concept identity is no longer decodable. The present findings provide preliminary evidence that this is not the case. Despite substantial heterogeneity in lesion size, location, and language impairment, individuals with aphasia showed decoding accuracies that were comparable to, and in some cases exceeded, those of healthy adults. Moreover, the RSA results suggested a broadly similar representational profile across groups, with experiential features showing the most consistent correspondence with neural similarity structure even in the presence of left-hemisphere damage. This preservation suggests that core aspects of semantic representation remain at least partially intact following stroke, consistent with models of distributed conceptual knowledge in which semantic information is supported across a broad network of heteromodal regions (e.g., Fernandino et al., 2022). Regions within the distributed semantic network may be especially important in this regard, given their established roles in multimodal semantic representation (Binder et al., 2009). Together, these findings are consistent with the view that impaired semantic access at the behavioral level in post-stroke aphasia does not necessarily reflect degraded conceptual representations and may instead reflect disrupted access to, or less efficient engagement of, a partially preserved semantic system.

Although concept-level semantic information remained decodable in participants with aphasia, the anatomical distribution of the most stable voxels provided important context for the regions contributing to decoding. In participants with aphasia, the largest voxel counts were observed in left middle occipital, left middle temporal, left precuneus, right angular, and left fusiform regions, with additional voxels distributed across bilateral temporal, occipital, and inferior parietal regions. Healthy participants showed a broadly overlapping distribution, with the largest voxel counts observed in left middle temporal, left fusiform, left middle occipital, left precuneus, and left parahippocampal regions, and additional voxels distributed across bilateral temporal, occipital, and inferior parietal regions. Together, these findings suggest that successful decoding in aphasia was supported by voxels distributed across a broad set of cortical regions that partially overlapped with the distribution observed in healthy adults. Although these anatomical distributions help contextualize the voxel sets supporting decoding, they should not be interpreted as direct evidence for functional reorganization. Rather, they more likely indicate that concept-level semantic information remained decodable in aphasia across stable voxels distributed within multiple spared cortical regions, consistent with the possibility that semantic information is supported by a distributed and partially redundant neural system.

The convergence of RSA, decoding, and voxel-localization findings provides insight into long-standing debates regarding the architecture of semantic representation. Across both individuals with aphasia and healthy adults, experiential features showed the most consistent correspondence with neural similarity structure, including after controlling for distributional and taxonomic models. This pattern aligns with distributed, grounded accounts of conceptual knowledge, which propose that semantic representations emerge from the convergence of perceptual, motor, affective, and contextual systems rather than from a single modality-independent hub (Barsalou, 2009; Binder et al., 2009; Lambon Ralph et al., 2017). The observation that Exp48 showed the most consistent correspondence with neural geometry in both groups, despite anatomical disruption in the aphasia group, suggests that feature-based semantic organization may remain an organizing principle even when portions of the left-hemisphere semantic network are damaged.

One important consideration is that the feature-generation task itself may have influenced which semantic model showed the strongest correspondence with neural data. Because participants were instructed to generate semantic features, the task may have preferentially engaged perceptual, motor, functional, and contextual attributes of concepts, thereby favoring an experiential model such as Exp48 over distributional or taxonomic models. However, the present findings are not unique to feature generation. Fernandino et al. (2022) similarly found that experiential features showed the strongest correspondence with neural similarity structure using RSA during a semantic familiarity task. Thus, although task demands may have increased the likelihood of participants generating experience-based features, converging evidence suggests that experiential dimensions capture a robust component of neural semantic organization across different semantic tasks.

Finally, the preserved decodability of conceptual information in aphasia, across stable voxels distributed in multiple spared cortical regions, suggests that concept-level semantic information is not reliant on a single locus or module. Instead, the present findings are consistent with a model in which conceptual information can be supported across multiple cortical regions. This pattern is consistent with the possibility that semantic representations are supported by a distributed and partially redundant neural system, although the present data do not directly establish redundancy or functional reorganization. Such distributed organization may contribute to the resilience of semantic representations following stroke, even when portions of the left-hemisphere semantic network are compromised. Together, these findings are consistent with a distributed, experience-based architecture of semantic representation, while providing less support for strictly modular accounts.

It is noteworthy that decoding accuracy was at chance level for control participants 006, 007, and 008. Although we cannot definitively determine why this pattern emerged, the RSA results provide some insight (see Figure 6). For these participants, there was no clear model that correlated most strongly with the BOLD signal. Moreover, participants 006 and 007 showed little to no alignment for any model. Because the semantic models sampled a broad range of proposed organizational principles for conceptual semantics, the absence of alignment across models is less easily explained by reliance on a single alternative semantic coding scheme not captured by the models tested here. One possible interpretation is that these participants were not consistently performing the semantic feature-generation task. A second possibility is that participants generated different features across trials for each item, which may have changed the BOLD signal associated with those items across runs. Although we hypothesized that the computational models of semantic representation chosen for this study would be resilient to this process, this possibility cannot be definitively ruled out.

Unexpectedly, behavioral measures of semantic processing, including Pyramids and Palm Trees, Camels and Cactus and s-weight, were not significantly associated with decoding accuracy. One possible interpretation is that overt semantic judgment performance depends on additional processes, such as cognitive control, response selection, and executive function, that are only partially related to the underlying representational geometry captured by the covert feature-generation task. However, these analyses were underpowered and vulnerable to outlier influence given the small aphasia sample. In contrast, the strong association between decoder accuracy and neural reliability suggests that decoding performance was more closely tied to the stability of the neural data than to the available behavioral measures. These findings are consistent with the possibility that semantic representations remained sufficiently intact to support decoding, even when behavioral access to those representations was more variable.

An important implication of the present findings is that decoding performance appears to have been shaped by the reliability and quantity of the neural data in addition to linguistic or clinical characteristics. Across all participants, decoder accuracy was strongly associated with split-half neural reliability, indicating that participants with more internally consistent neural representational geometry achieved higher decoding performance. Decoder accuracy was also positively associated with the number of functional runs completed, suggesting that dense sampling improved the stability of item-level beta estimates and, in turn, the fidelity of the neural similarity structure used for decoding.

Although the present study was not designed to evaluate a treatment protocol, the results provide a useful bridge to clinical approaches such as SFA, which explicitly targets feature generation as a means of strengthening lexical-semantic mappings. The covert feature-generation task used in our fMRI paradigm engages the same underlying operation central to SFA, accessing and assembling semantic features of a concept, yet does so in a context that isolates representational processing from overt speech demands. Thus, the neural representational structures captured here offer a window into the integrity of conceptual knowledge as it is accessed during a process analogous to one of the active ingredients of SFA.

The finding that experiential semantic similarity showed the most consistent correspondence with neural representational geometry in participants with aphasia and healthy adults aligns with theoretical accounts proposing that feature-based treatments such as SFA may draw on richly grounded conceptual features. Prior behavioral work has shown that producing more semantically proximal features is associated with better naming outcomes (Cavanaugh et al., 2026). The present results extend this logic to the neural domain: during covert feature generation, the representational space most consistently aligned with neural similarity structure was the one organized around experiential attributes. However, this interpretation should be considered in light of the task itself. Because participants were explicitly instructed to generate semantic features, the task may have preferentially engaged sensory, motor, functional, and contextual properties of concepts, thereby favoring alignment with an experiential model such as Exp48 over distributional or taxonomic models. Thus, the present findings suggest that experiential semantic structure remains accessible and neurally measurable during feature generation in chronic aphasia, while also highlighting that different tasks may evoke different semantic geometries.

At the same time, the RSD findings highlight important constraints on how semantic information is accessed in aphasia. Although conceptual representations remained decodable, the stable voxels supporting decoding in participants with aphasia were distributed across spared cortical regions, suggesting that concept-level semantic information can remain measurable across distributed cortical tissue after stroke. Together, these findings suggest that future work on SFA and similar feature-based treatments (Sandberg et al., 2023) may benefit from incorporating measures that assess not only behavioral output but also the stability of semantic access processes. The present study provides preliminary neural evidence that the conceptual-semantic representations engaged by feature generation remain decodable in chronic aphasia, supporting the possibility that treatment may leverage partially preserved representational content while working to improve access and control mechanisms.

Several limitations should be considered when interpreting the present findings. First, the sample size was modest for both groups. Although dense sampling substantially increased within-participant reliability, the small number of participants limits power for detecting individual-differences relationships and constrains the generalizability of group-level effects. For example, the absence of statistically significant correlations between decoding accuracy and behavioral semantic measures should be interpreted cautiously; larger samples are required to determine whether these null effects reflect true dissociations or insufficient statistical power.

Second, the degree of lesion heterogeneity introduces variability that is both clinically representative and analytically challenging. Participants differed in lesion size, cortical extent, and involvement of classical semantic regions, making it difficult to isolate lesion–symptom relationships or define clear anatomical predictors of decoding performance. A larger, anatomically stratified cohort would allow stronger inferences about how specific regions contribute to representational integrity and altered voxel distributions following stroke-induced aphasia.

Third, although the covert semantic feature generation task effectively engages conceptual retrieval while minimizing speech demands, it does not capture overt feature production or control processes that may shape performance in therapeutic contexts such as Semantic Feature Analysis. Covert tasks reduce motor-related confounds but may underestimate the role of executive or control-based retrieval demands that influence functional communication. Furthermore, they may also reduce task engagement and do not permit experimenters to assess the validity of participant responses.

Fourth, although we controlled for visual similarity using a visual RDM derived from ResNet-50, this procedure cannot fully eliminate all visual confounds. Residual perceptual structure may still contribute subtly to neural similarity patterns within higher visual or ventral temporal regions.

A critical next step for this line of research is to extend representational analyses beyond single-word semantic processing to the sentence- and discourse-level communication demands that matter most to people with aphasia. Recent consensus work emphasizes that the priorities of individuals living with aphasia center on everyday communication, including conversational participation, functional comprehension, and successful interaction with communication partners, rather than gains limited to word-level naming or feature retrieval tasks (Shrubsole et al., 2024). It is possible that moving beyond single words may also change the neural representational geometry, for example, to be more consistent with distributional or taxonomic models. This perspective highlights an important implication of the present findings: although core conceptual-semantic representations remained decodable at the single-word level, successful communication depends on integrating semantic information across time, sentences, discourse contexts, and interactional settings.

Building on the present word-level decoding results, future work should examine the accessibility and neural organization of semantic information during connected speech, sentence comprehension, and narrative feature generation. Extending RSA and RSD frameworks to naturalistic or semi-naturalistic tasks, such as sentence-level feature generation, story retell, or conversational turn-taking would allow us to test whether the preserved experiential representational geometry observed here generalizes to richer linguistic contexts. Such work would better align neural representational analysis with the functional communication goals prioritized by stakeholders. It would also clarify how representational integrity interacts with dynamic processes such as semantic integration, predictive processing, and conceptual combination, all of which are essential for everyday communication but remain underexamined in neuroimaging studies of aphasia. In this way, future research can advance both the scientific understanding of semantic resilience and the clinical goal of improving meaningful communication participation for people with aphasia.

## CRediT author statement

Swiderski, Alexander: Conceptualization (lead), Methodology (lead), Software (lead), Validation (lead), Formal analysis (lead), Resources (equal), Data Curation (equal), Writing - Original Draft (lead), Writing - Review & Editing (equal), Visualization (lead), Supervision (equal), Project administration (equal), Funding acquisition (Lead).

Bohland, Jason: Conceptualization (supporting), Methodology (supporting), Validation (supporting), Formal analysis (supporting), Data Curation (supporting), Writing - Review & Editing (supporting), Visualization (supporting), Supervision (supporting), and Funding acquisition (supporting).

Dickey, Michael Walsh: Conceptualization (supporting), Methodology (supporting), Validation (supporting), Formal analysis (supporting), Resources (supporting), Writing - Original Draft (supporting), Writing - Review & Editing (supporting), Supervision (equal), Project administration (equal), and Funding acquisition (supporting).

Johnson, Jeffrey P.: Methodology (supporting), Software (supporting), Validation (supporting), Writing - Review & Editing (supporting), Visualization (supporting), and Funding acquisition (supporting).

Wilson, Stephen M: Methodology (supporting), Software (supporting), Validation (supporting), Writing - Review & Editing (supporting), Visualization (supporting), and Funding acquisition (supporting).

Hula, William D: Conceptualization (supporting), Methodology (supporting), Software (supporting), Validation (supporting), Formal analysis (supporting), Resources (supporting), Data Curation (supporting), Writing - Original Draft (supporting), Writing - Review & Editing (supporting), Visualization (supporting), Supervision (supporting), Project administration (supporting), Funding acquisition (supporting).

## Acknowledgements

We are sincerely thankful for study participants for their time and commitment to this study. We are also thankful for Mackenna Locke who developed the training materials for this study.

