## supplemental tables for "Concept-Level Semantic Representations Remain Decodable in Chronic Post-Stroke Aphasia"

Supplemental Table 1. Participants with aphasia: Lesioned voxel count within the Semantic Network ROI partitioned by the Automated Anatomical Labeling Atlas

|  | sub-101 | sub-102 | sub-103 | sub-105 | sub-106 | sub-107 | group sum | group mean |
| --- | --- | --- | --- | --- | --- | --- | --- | --- |
| Left Middle Occipital | 59 | 31 | 106 | 45 | 142 | 146 | 529 | 151.14 |
| Right Middle Occipital | 41 | 20 | 12 | 2 | 77 | 21 | 173 | 49.43 |
| Left Fusiform | 4 | 73 | 69 | 21 | 1 | 5 | 173 | 49.43 |
| Left Angular | 9 | 8 | 32 | 52 | 8 | 25 | 134 | 38.29 |
| Left Middle Temporal | 55 | 99 | 9 | 79 | 164 | 10 | 416 | 118.86 |
| Right Middle Temporal | 6 | 10 | 1 | 1 | 1 | 39 | 58 | 16.57 |
| Left Inferior Temporal | 10 | 21 | 8 | 1 | 2 | 1 | 43 | 12.29 |
| Left Inferior Frontal Triangularis | 8 | 25 | 16 | 17 | 0 | 1 | 67 | 19.14 |
| Left Inferior Parietal | 12 | 25 | 0 | 90 | 4 | 1 | 132 | 37.71 |
| Right Angular | 34 | 13 | 0 | 2 | 89 | 74 | 212 | 60.57 |
| Left Precuneus | 80 | 20 | 68 | 47 | 0 | 6 | 221 | 63.14 |
| Right Precuneus | 50 | 16 | 43 | 19 | 0 | 37 | 165 | 47.14 |
| Right Inferior Temporal | 0 | 28 | 27 | 17 | 10 | 6 | 88 | 25.14 |
| Left Inferior Frontal Opercularis | 1 | 27 | 0 | 23 | 0 | 4 | 55 | 15.71 |
| Left Parahippocampal | 2 | 9 | 24 | 0 | 0 | 1 | 36 | 10.29 |
| Right Calcarine | 16 | 0 | 20 | 0 | 0 | 6 | 42 | 12.00 |
| Left Cuneus | 3 | 3 | 0 | 0 | 0 | 4 | 10 | 2.86 |
| Right Lingual | 9 | 0 | 8 | 0 | 0 | 5 | 22 | 6.29 |
| Left Supramarginal | 0 | 6 | 0 | 6 | 0 | 12 | 24 | 6.86 |
| Right Posterior Cingulum | 0 | 0 | 1 | 2 | 0 | 3 | 6 | 1.71 |
| Left Superior Frontal | 20 | 20 | 0 | 0 | 0 | 0 | 40 | 11.43 |
| Left Middle Frontal | 8 | 0 | 17 | 0 | 0 | 0 | 25 | 7.14 |
| Left Inferior Frontal Orbital | 10 | 0 | 0 | 4 | 0 | 0 | 14 | 4.00 |
| Left Anterior Cingulum | 4 | 0 | 0 | 1 | 0 | 0 | 5 | 1.43 |
| Left Middle Cingulum | 4 | 0 | 10 | 0 | 0 | 0 | 14 | 4.00 |
| Left Calcarine | 9 | 0 | 3 | 0 | 0 | 0 | 12 | 3.43 |
| Left Superior Temporal Pole | 2 | 0 | 0 | 11 | 0 | 0 | 13 | 3.71 |
| Left Olfactory | 0 | 1 | 0 | 0 | 0 | 2 | 3 | 0.86 |
| Left Insula | 0 | 5 | 0 | 0 | 0 | 32 | 37 | 10.57 |
| Left Posterior Cingulum | 0 | 8 | 0 | 14 | 0 | 0 | 22 | 6.29 |
| Left Superior Occipital | 0 | 3 | 3 | 0 | 0 | 0 | 6 | 1.71 |
| Right Superior Temporal | 0 | 1 | 0 | 0 | 0 | 8 | 9 | 2.57 |
| Left Rectus | 0 | 0 | 1 | 3 | 0 | 0 | 4 | 1.14 |
| Right Middle Frontal | 0 | 0 | 0 | 6 | 0 | 2 | 8 | 2.29 |
| Left Medial Frontal Orbital | 0 | 0 | 0 | 1 | 0 | 2 | 3 | 0.86 |
| Right Middle Cingulum | 0 | 0 | 0 | 1 | 0 | 1 | 2 | 0.57 |
| Left Precentral | 2 | 0 | 0 | 0 | 0 | 0 | 2 | 0.57 |
| Left Superior Medial Frontal | 15 | 0 | 0 | 0 | 0 | 0 | 15 | 4.29 |
| Right Insula | 1 | 0 | 0 | 0 | 0 | 0 | 1 | 0.29 |
| Left Hippocampus | 8 | 0 | 0 | 0 | 0 | 0 | 8 | 2.29 |
| Left Thalamus | 2 | 0 | 0 | 0 | 0 | 0 | 2 | 0.57 |
| Left Superior Temporal | 1 | 0 | 0 | 0 | 0 | 0 | 1 | 0.29 |
| Right Fusiform | 0 | 7 | 0 | 0 | 0 | 0 | 7 | 2.00 |
| Right Medial Frontal Orbital | 0 | 0 | 0 | 3 | 0 | 0 | 3 | 0.86 |
| Left Superior Parietal | 0 | 0 | 0 | 14 | 0 | 0 | 14 | 4.00 |
| Other | 15 | 21 | 22 | 18 | 2 | 46 | 124 | 20.67 |

Supplemental Table 2. Control participants: Voxel count by region location within the Automated Anatomical Labeling Atlas

|  | sub-001 | sub-002 | sub-003 | sub-004 | sub-005 | sub-006 | sub-007 | sub-008 | group sum | group mean |
| --- | --- | --- | --- | --- | --- | --- | --- | --- | --- | --- |
| Left Middle Temporal | 122 | 56 | 123 | 7 | 248 | 30 | 37 | 20 | 643 | 80.375 |
| Left Fusiform | 12 | 36 | 121 | 8 | 63 | 40 | 8 | 2 | 290 | 36.25 |
| Left Middle Occipital | 26 | 71 | 40 | 20 | 37 | 36 | 3 | 42 | 275 | 34.375 |
| Left Parahippocampal | 5 | 45 | 63 | 3 | 24 | 50 | 2 | 6 | 198 | 24.75 |
| Left Inferior Temporal | 0 | 3 | 7 | 6 | 5 | 3 | 2 | 1 | 27 | 3.375 |
| Right Middle Temporal | 22 | 8 | 39 | 0 | 79 | 0 | 48 | 1 | 197 | 24.625 |
| Left Inferior Parietal | 8 | 70 | 0 | 44 | 0 | 11 | 1 | 37 | 171 | 21.375 |
| Right Precuneus | 33 | 48 | 37 | 28 | 0 | 1 | 19 | 0 | 166 | 20.75 |
| Left Cuneus | 31 | 13 | 1 | 27 | 0 | 1 | 0 | 1 | 74 | 9.25 |
| Left Precuneus | 56 | 4 | 0 | 123 | 0 | 4 | 12 | 0 | 199 | 24.875 |
| Right Inferior Temporal | 24 | 25 | 28 | 0 | 18 | 14 | 0 | 0 | 109 | 13.625 |
| Left Angular | 49 | 0 | 0 | 16 | 10 | 0 | 24 | 4 | 103 | 12.875 |
| Left Calcarine | 50 | 2 | 0 | 36 | 0 | 6 | 1 | 0 | 95 | 11.875 |
| Right Angular | 21 | 0 | 2 | 6 | 0 | 2 | 5 | 0 | 36 | 4.5 |
| Left Middle Frontal | 0 | 0 | 0 | 1 | 0 | 34 | 71 | 49 | 155 | 19.375 |
| Right Middle Occipital | 0 | 51 | 11 | 0 | 0 | 25 | 0 | 47 | 134 | 16.75 |
| Left Superior Frontal | 0 | 0 | 0 | 7 | 0 | 2 | 29 | 89 | 127 | 15.875 |
| Left Inferior Frontal Orbital | 0 | 0 | 0 | 3 | 0 | 14 | 49 | 45 | 111 | 13.875 |
| Left Inferior Frontal Triangularis | 0 | 0 | 0 | 3 | 0 | 81 | 10 | 16 | 110 | 13.75 |
| Left Superior Medial Frontal | 0 | 0 | 0 | 8 | 0 | 1 | 27 | 68 | 104 | 13 |
| Left Inferior Frontal Opercularis | 0 | 0 | 0 | 7 | 0 | 42 | 6 | 26 | 81 | 10.125 |
| Left Insula | 0 | 0 | 0 | 9 | 0 | 53 | 17 | 1 | 80 | 10 |
| Right Calcarine | 15 | 20 | 5 | 4 | 0 | 0 | 0 | 0 | 44 | 5.5 |
| Right Superior Temporal | 6 | 0 | 2 | 0 | 0 | 8 | 0 | 1 | 17 | 2.125 |
| Left Superior Temporal | 0 | 0 | 0 | 0 | 2 | 2 | 7 | 2 | 13 | 1.625 |
| Right Lingual | 13 | 25 | 0 | 23 | 0 | 0 | 0 | 0 | 61 | 7.625 |
| Left Supramarginal | 0 | 0 | 0 | 1 | 0 | 5 | 0 | 34 | 40 | 5 |
| Left Anterior Cingulum | 0 | 0 | 0 | 17 | 0 | 0 | 18 | 1 | 36 | 4.5 |
| Left Superior Temporal Pole | 0 | 0 | 0 | 13 | 0 | 1 | 13 | 0 | 27 | 3.375 |
| Left Medial Frontal Orbital | 0 | 0 | 0 | 3 | 0 | 0 | 14 | 2 | 19 | 2.375 |
| Left Superior Parietal | 0 | 14 | 0 | 2 | 0 | 1 | 0 | 0 | 17 | 2.125 |
| Left Middle Cingulum | 0 | 0 | 0 | 10 | 0 | 0 | 1 | 1 | 12 | 1.5 |
| Right Fusiform | 0 | 0 | 5 | 0 | 2 | 1 | 0 | 0 | 8 | 1 |
| Right Parahippocampal | 0 | 0 | 4 | 0 | 1 | 0 | 0 | 1 | 6 | 0.75 |
| Left Posterior Cingulum | 0 | 0 | 0 | 24 | 0 | 0 | 7 | 0 | 31 | 3.875 |
| Right Insula | 0 | 0 | 0 | 8 | 0 | 0 | 5 | 0 | 13 | 1.625 |
| Right Inferior Frontal Opercularis | 0 | 0 | 0 | 2 | 0 | 0 | 3 | 0 | 5 | 0.625 |
| Right Middle Frontal | 0 | 0 | 0 | 3 | 0 | 0 | 1 | 0 | 4 | 0.5 |
| Right Middle Cingulum | 0 | 0 | 0 | 1 | 0 | 3 | 0 | 0 | 4 | 0.5 |
| Right Posterior Cingulum | 0 | 0 | 0 | 3 | 0 | 0 | 1 | 0 | 4 | 0.5 |
| Left Hippocampus | 0 | 0 | 0 | 0 | 0 | 2 | 2 | 0 | 4 | 0.5 |
| Right Medial Frontal Orbital | 0 | 0 | 0 | 2 | 0 | 0 | 1 | 0 | 3 | 0.375 |
| Left Rectus | 1 | 0 | 1 | 0 | 0 | 0 | 0 | 0 | 2 | 0.25 |
| Right Inferior Frontal Triangularis | 0 | 0 | 0 | 3 | 0 | 0 | 0 | 0 | 3 | 0.375 |
| Left Superior Occipital | 0 | 0 | 0 | 0 | 0 | 3 | 0 | 0 | 3 | 0.375 |
| Left Olfactory | 0 | 0 | 0 | 0 | 0 | 0 | 2 | 0 | 2 | 0.25 |
| Left Thalamus | 0 | 0 | 0 | 0 | 0 | 0 | 2 | 0 | 2 | 0.25 |
| Right Middle Temporal Pole | 0 | 0 | 0 | 0 | 0 | 0 | 0 | 2 | 2 | 0.25 |
| Right Superior Temporal Pole | 0 | 0 | 0 | 0 | 0 | 0 | 1 | 0 | 1 | 0.125 |
| Right Hippocampus | 0 | 0 | 0 | 0 | 0 | 0 | 0 | 1 | 1 | 0.125 |
| Other | 6 | 9 | 11 | 19 | 11 | 24 | 51 | 0 | 131 | 16.375 |
